# Modulation of sensory behavior and food choice by an enteric bacteria-produced neurotransmitter

**DOI:** 10.1101/735845

**Authors:** Michael P. O’Donnell, Bennett W. Fox, Pin-Hao Chao, Frank C. Schroeder, Piali Sengupta

## Abstract

Animals coexist in commensal, pathogenic or mutualistic relationships with complex communities of diverse organisms including microbes^1^. Some bacteria produce bioactive neurotransmitters which have been proposed to modulate host nervous system activity and behaviors^2^. However, the mechanistic basis of this microbiota-brain modulation and its physiological relevance is largely unknown. Here we show that in *C. elegans*, the neuromodulator tyramine (TA) produced by gut-colonizing commensal *Providencia* bacteria can bypass the requirement for host TA biosynthesis to manipulate a host sensory decision. Bacterially-produced TA is likely converted to octopamine (OA) by the host tyramine beta-hydroxylase enzyme. OA, in turn, targets the OCTR-1 receptor on the ASH/ASI sensory neurons to modulate an aversive olfactory response. We identify genes required for TA biosynthesis in *Providencia*, and show that these genes are necessary for modulation of host behavior. We further find that *C. elegans* colonized by *Providencia* preferentially select these bacteria in food choice assays, and that this selection bias requires bacterially-produced TA. Our results demonstrate that a neurotransmitter produced by gut microbiota mimics the functions of the cognate host molecule to override host control of a sensory decision, thereby promoting fitness of both host and microbe.

---

The pathways mediating chemical communication between gut-colonizing bacteria and the host nervous system are largely undescribed^2^. Recently, the nematode *C. elegans* has emerged as a powerful system in which to study host-microbe chemical communication^3^, offering an opportunity to experimentally address how microbiota influence host nervous system function. Diverse populations of pathogenic and non-pathogenic bacteria both colonize the *C. elegans* intestine and serve as its primary food source in the wild^4^. Exposure to pathogenic bacteria alters *C. elegans* olfactory behaviors^5^, but whether commensal gut bacteria also modulate host behaviors is unknown^4^.

To identify novel modes of microbial influences on host sensory behaviors, we screened non-pathogenic bacterial strains typically associated with wild nematodes^6^ for their ability to influence *C. elegans* olfactory responses. In long-range chemotaxis assays^7^, adult hermaphrodites co-cultivated on these bacterial strains exhibited robust attraction to a panel of attractive volatile odorants similar to the behaviors of animals grown on the standard *E. coli* food source OP_50_ (Fig. 1a). However, co-cultivation with the *Providencia alcalifaciens* strain (JUb_39_)^6,8^ resulted in decreased avoidance of 100% 1-octanol as compared to OP_50_-grown animals (Fig. 1b; this decreased avoidance is henceforth referred to as octanol modulation). Avoidance of the volatile and osmotic repellents 2-nonanone and 8M glycerol, respectively, was unaffected upon growth on JUb_39_ (Fig. 1b, Fig. S1a), suggesting that JUb_39_ modulates responses of *C. elegans* to a selective subset of nociceptive chemical stimuli. Animals grown on a distantly-related *Providencia rettgeri* strain isolated from nematodes in compost (PYb_007_, Fig. S1b) exhibited similar octanol modulation (Fig. 1c). These observations indicate that upon co-culture, multiple *Providencia* strains modulate octanol aversion in *C. elegans*.

**Fig. 1.**
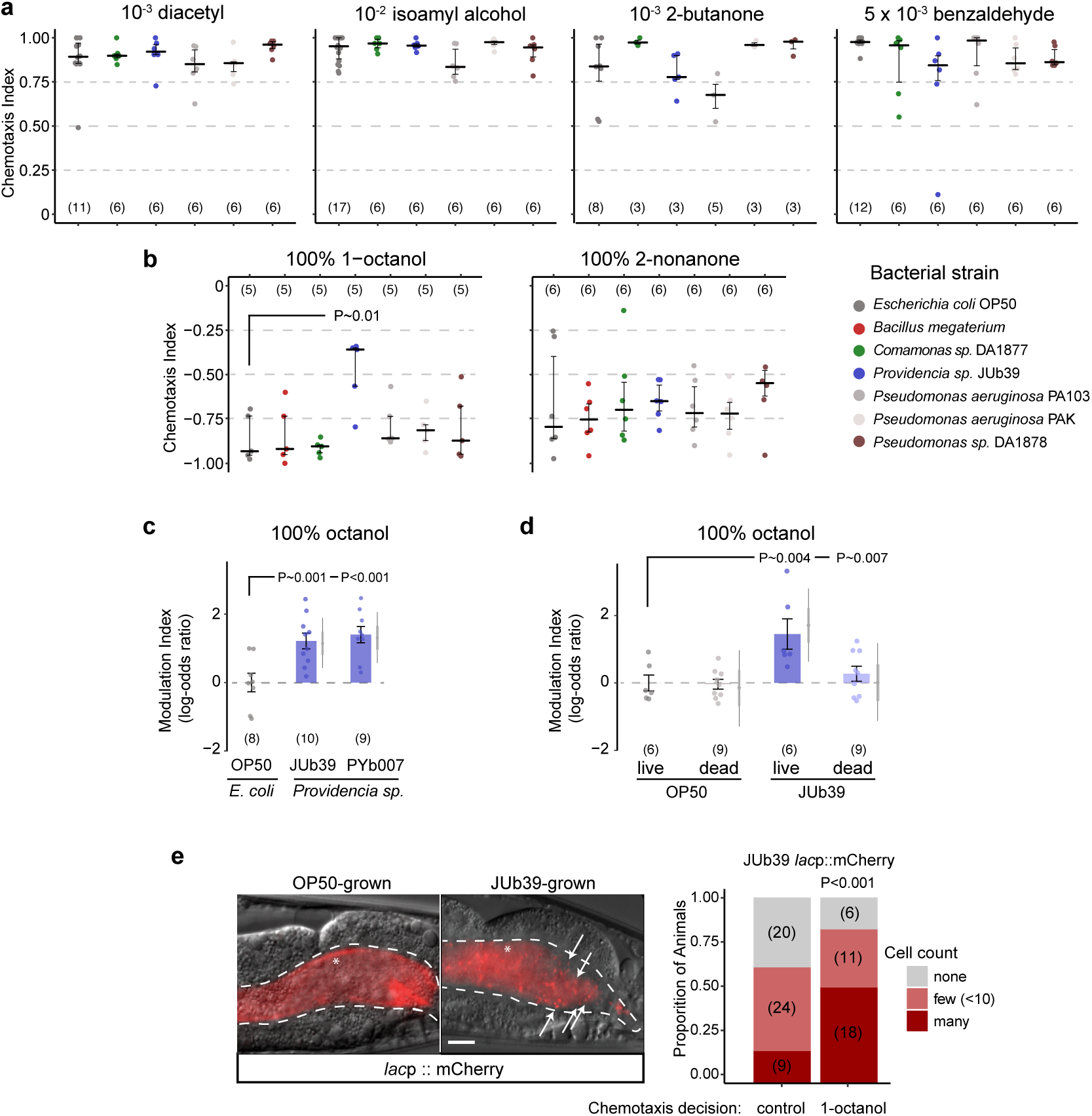
*Providencia* colonizes the *C. elegans* intestine and modulates octanol avoidance behavior. **a-b)** Long-range chemotaxis assays of *C. elegans* grown on the indicated bacterial strains to attractive **(a)** or aversive **(b)** odors. Chemotaxis index (CI) = (animals at the odorant – animals at the control)/total number of animals. Each dot indicates the CI from a single assay of approximately 100 animals. Positive and negative numbers indicate attraction and avoidance, respectively. Horizontal line is median; errors are 1st and 3rd quartiles. *P*-value indicated is from a binomial general linearized mixed-effects model (GLMM) with random intercepts for assay plate and date and with false discovery rate (FDR) for post-hoc comparisons. Numbers in parentheses indicate total number of assays. **c-d)** Modulation index of worms grown on the indicated bacterial strains **(c)** or bacterial strains pre-treated with 200 *µ*g/*µ*L gentamicin for 2 hrs prior to plating **(d)** in response to 100% octanol. Modulation index is defined as the log odds-ratio of the proportion of worms at octanol vs control of each condition relative to the OP_50_-grown condition per independent day. Modulation index values are shown on a log-odds (logit) scale and are normalized to the values of wild-type animals grown on OP_50_ for each day, indicated with a gray dashed line. Positive numbers indicate reduced octanol avoidance. Errors are SEM. Gray thin and thick vertical bars at right indicate Bayesian 95% and 66% credible intervals, respectively. *P*-values between the indicated conditions are from a GLMM with Dunnett-type multivariate-t adjustment for **c**, and Tukey-type multivariate-t adjustment for **d**. **e)** Presence of mCherry-expressing bacteria in the posterior intestines of young adult animals indicated with micrographs (left) or quantified (right). Arrows in micrographs indicate intact rod-shaped cells, asterisk indicates diffuse intestinal fluorescence. Dashed line in micrographs indicate the intestinal boundary. Anterior is at left. Scale bar: 10 *µ*m. Bars at right show proportion of animals with the indicated distribution of JUb_39_ cells present in animals that migrated to 100% octanol or the control in chemotaxis assays. Numbers in parentheses indicate the number of animals; 3 independent assays. *P*-value is derived from an ordinal regression.

Under specific conditions, food deprivation reduces octanol avoidance^9^. JUb_39_ has been categorized as a ‘beneficial’ bacterium that supports robust *C. elegans* growth and does not induce stress responses^6^, suggesting that JUb_39_-fed animals are unlikely to be nutrition-deprived. In support of this notion, growth on JUb_39_ did not alter expression of a *tph-1p::gfp* fusion gene, a reporter of feeding state^10,11^ (Fig. S1c). Moreover, growth of *C. elegans* on the poor bacterial food *Bacillus megaterium*^12^ did not alter octanol avoidance (Fig. 1b). We infer that the observed octanol modulation by *Providencia* is unlikely to be solely due to changes in feeding state.

While OP_50_ is typically crushed by the pharyngeal grinder in young adult *C. elegans*^13^, a subset of bacterial strains can bypass the grinder and survive in the worm intestine^6,14,15^. We found that feeding *C. elegans* with JUb_39_ pre-treated with high concentrations of the antibiotic gentamicin eliminated octanol modulation (Fig. 1d), indicating that JUb_39_ must be alive to mediate this behavioral plasticity. In addition, neither exposure of OP_50_-grown animals to JUb_39_-derived odors nor pre-incubation of OP_50_-grown animals with JUb_39_-conditioned media was sufficient to result in octanol modulation (Fig. S1d-e), further suggesting that *C. elegans* must ingest live JUb_39_ to induce octanol modulation.

To test whether colonization of the worm gut drives octanol modulation, we transformed OP_50_ and JUb_39_ with a plasmid encoding a constitutively expressed mCherry fluorescent reporter. While the guts of OP_50_-fed adult worms displayed only diffuse intestinal fluorescence consistent with these bacteria being lysed, the guts of JUb_39_-fed worms contained variable but typically large numbers of intact rod-shaped cells expressing mCherry (Fig. 1e), likely indicating the presence of live JUb_39_. These cells tended to be enriched in the posterior intestine (Fig. 1e), unlike the reported localization pattern of severely pathogenic bacteria^16^. Moreover, nematodes colonized by JUb_39_ did not exhibit phenotypes characteristic of pathogenic infection such as anal swelling^17^ (Fig. 1e), further confirming that JUb_39_ is largely non-pathogenic to *C. elegans*.

We next performed chemotaxis assays with animals fed on mCherry-labeled JUb_39_, and quantified intestinal bacterial cells in animals that had navigated either toward octanol or toward the control. We found that animals navigating toward octanol consistently contained more gut bacteria (Fig. 1e). We conclude that JUb_39_ colonizes the worm gut and the extent of colonization is correlated with decision-making in response to octanol.

We investigated the mechanistic basis for *Providencia*-mediated octanol modulation. Octanol avoidance is subject to extensive modulation directly and indirectly via multiple biogenic amines including tyramine (TA) and octopamine (OA) (Fig. 2a) as well as neuropeptides^9,18-22^. TA is produced from Tyrosine (L-Tyr) via the activity of a tyrosine decarboxylase (TDC; encoded by *tdc-1* in *C. elegans*); TA is subsequently converted to OA via a tyramine beta hydroxylase (encoded by *tbh-1*)^23^ (Fig. 2a). Consequently, all *tbh-1* mutant phenotypes resulting from lack of OA are expected to be shared by *tdc-1* mutants^23^. Unexpectedly, we found that while *tdc-1* mutants grown on JUb_39_ continued to exhibit octanol modulation, the modulation exhibited by *tbh-1* mutants was significantly reduced (Fig. 2b). Mutations in the *cat-2* tyrosine hydroxylase^24^ and *tph-1* tryptophan hydroxylase^25^ enzymes required for the production of biogenic amines dopamine and serotonin in *C. elegans*, respectively, did not affect octanol modulation (Fig. 2b). These results raise the possibility that *C. elegans*-produced OA, but not TA, is partly necessary for JUb_39_-mediated octanol modulation.

**Fig. 2.**
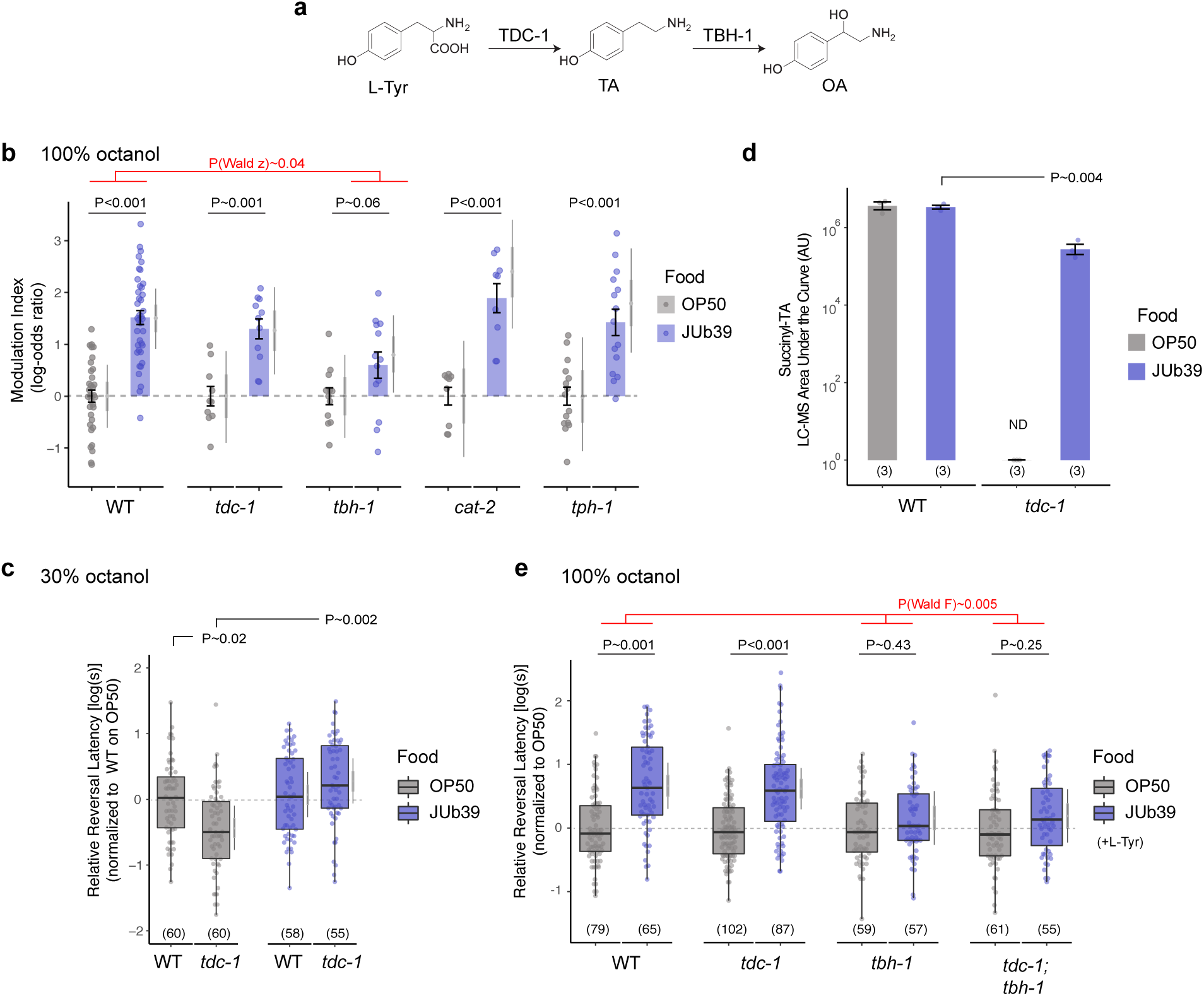
*Providencia* produces TA which compensates for loss of *C. elegans tdc-1*. **a)** Biosynthesis pathway of TA and OA from L-Tyr in *C. elegans*^23^. **b)** Modulation index of worms of the indicated genotypes grown on OP_50_ or JUb_39_ in response to 100% octanol. Each dot represents results from one chemotaxis assay with approximately 100 animals each. Values are shown on the log-odds (logit) scale and are normalized to the values of animals of the corresponding genotypes grown on OP_50_ for each day, indicated with gray dashed lines. Positive numbers indicate reduced avoidance of octanol. Errors are SEM. Gray thin and thick vertical bars at right indicate Bayesian 95% and 66% credible intervals, respectively. *P*-values between the indicated conditions are from a GLMM with Dunnett-type multivariate-t adjustment. *P*-value in red indicates Wald z-statistic for the magnitude of the JUb_39_ effect in *tbh-1* compared to wild-type. **c, e)** Reversal response latency of animals of the indicated genotypes grown on OP_50_ or JUb_39_ on NGM **(c)** or NGM + 0.5% L-Tyr **(e)** in response to 30% octanol **(c)** or 100% octanol **(e)** in SOS assays. Each dot is the response time of a single animal. The Y-axis is log_10_-scaled for these log-normal distributed data, and normalized to the indicated control group for each experimental day. Numbers in parentheses indicate the number of worms tested in assays over at least 3 independent days. Boxplot indicates median and quartiles, whiskers indicate the data range, excluding outliers. Gray thin and thick vertical bars at right indicate Bayesian 95% and 66% credible intervals for the difference of means, respectively. *P*-values between the indicated conditions are from a linear-mixed effects regression on log-transformed data (LMM). *P*-value in red **(e)** indicates Wald F-statistic for the effect of the indicated genotypes on the magnitude of the JUb_39_ effect. **d)** Quantification of succinyl-TA in wild-type and *tdc-1* mutant animals grown on either OP_50_ or JUb_39_. Data are averaged from three independent replicates each. ND, not detected. *P*-values are from two-way ANOVA.

To account for these observations, we hypothesized that JUb_39_ may produce TA that functionally compensates for the host *tdc-1* mutation. *tdc-1* mutants grown on OP_50_ have been reported to display more rapid aversive responses to dilute (30%) octanol^18^. Exogenous TA suppresses this increased aversion of *tdc-1* animals but does not alter wild-type responses under the same conditions^18^. To test if JUb_39_ is able to suppress behavioral defects of *tdc-1* mutants, we performed short-range acute avoidance assays [the “smell-on-a-stick” (SOS) assay]^9,26^. In this assay, the strength of avoidance is inversely correlated with reversal latency when the animal encounters the repellent as it is moving forward. As expected, *tdc-1* mutants grown on OP_50_ responded more rapidly to 30% octanol than wild-type animals (Fig. 2c). This enhanced aversion was suppressed upon growth on JUb_39_ (Fig. 2c). These results are consistent with the notion that bacterially-produced TA functionally complements for the loss of host-derived TA in driving a sensory behavioral decision.

To directly test for the production of TA by JUb_39_, we measured succinyl-TA levels using high-resolution HPLC-MS in wild-type and *tdc-1* worms grown on either OP_50_ or JUb_39_^27^. Levels of succinyl-TA were comparable in wild-type animals grown on either OP_50_ or JUb_39_, whereas *tdc-1* mutants grown on OP_50_ had no detectable succinyl-TA, consistent with previous reports^23^ (Fig. 2d, Fig. S2). Notably, succinyl-TA levels were partly restored in *tdc-1* mutants grown on JUb_39_ (Fig. 2d). OA was not detected under these conditions in either wild type or *tdc-1* mutants. We conclude that JUb_39_ in association with *C. elegans* produces TA which can accumulate in the host.

Although TA biosynthesis in bacteria has been demonstrated in some gram-positive genera, production appears to be uncommon in gram-negative bacteria which include *Providencia*^28,29^. TA production in gram-positive strains is induced upon supplementation with L-Tyr^30^. We found that growth on L-Tyr-supplemented media enhanced octanol modulation by JUb_39_ in SOS assays (Fig. S3). Under these conditions, mutations in *tbh-1* fully suppressed octanol modulation in SOS assays, whereas consistent with our observations in long-range chemotaxis assays, *tdc-1* mutants continued to exhibit robust octanol modulation (Fig. 2e). Octanol avoidance behaviors of *tdc-1; tbh-1* double mutants were similar to those of *tbh-1* mutants alone (Fig. 2e), indicating that the lack of host-derived OA, and not accumulation of TA due to loss of TBH-1^23,27^ accounts for the reduced octanol modulation in *tbh-1* mutants.

Biogenic amines are typically generated from aromatic amino acids and L-glutamate by pyridoxyl phosphate (PLP)-dependent group II aromatic amino acid decarboxylase enzymes (AADCs) in both eukaryotes and bacteria^31^ (Fig. S4a-b, Table S1). In gram-positive *Enterococcus* and *Lactobacillus* (*Lb*), TA production is mediated by the TDC-encoding (*tyrDC*) AADC and tyrosine permease/transporter (*tyrP*) genes present in an operon; this operon is inducible by L-Tyr (Fig. 3a-b, S4a-b, Table S1)^32,33^. Although genes related to *Enterococcus tyrDC* and *tyrP* were largely absent in *Gammaproteobacteria* (Fig. 3b), we confirmed the presence of homologous operons containing *tyrDC* and *tyrP* in JUb_39_ and PYb_007_ in *de novo* genome assemblies via whole genome sequencing (Fig. 3a, Fig. S4b, Table S1). *tyrDC* homologs were also identified in the genomes of additional members of the *Morganellaceae* family, although the operon structure was conserved in only a subset of these genomes (Fig. 3a, Fig. 3c, S4b, Table S1).

**Fig. 3.**
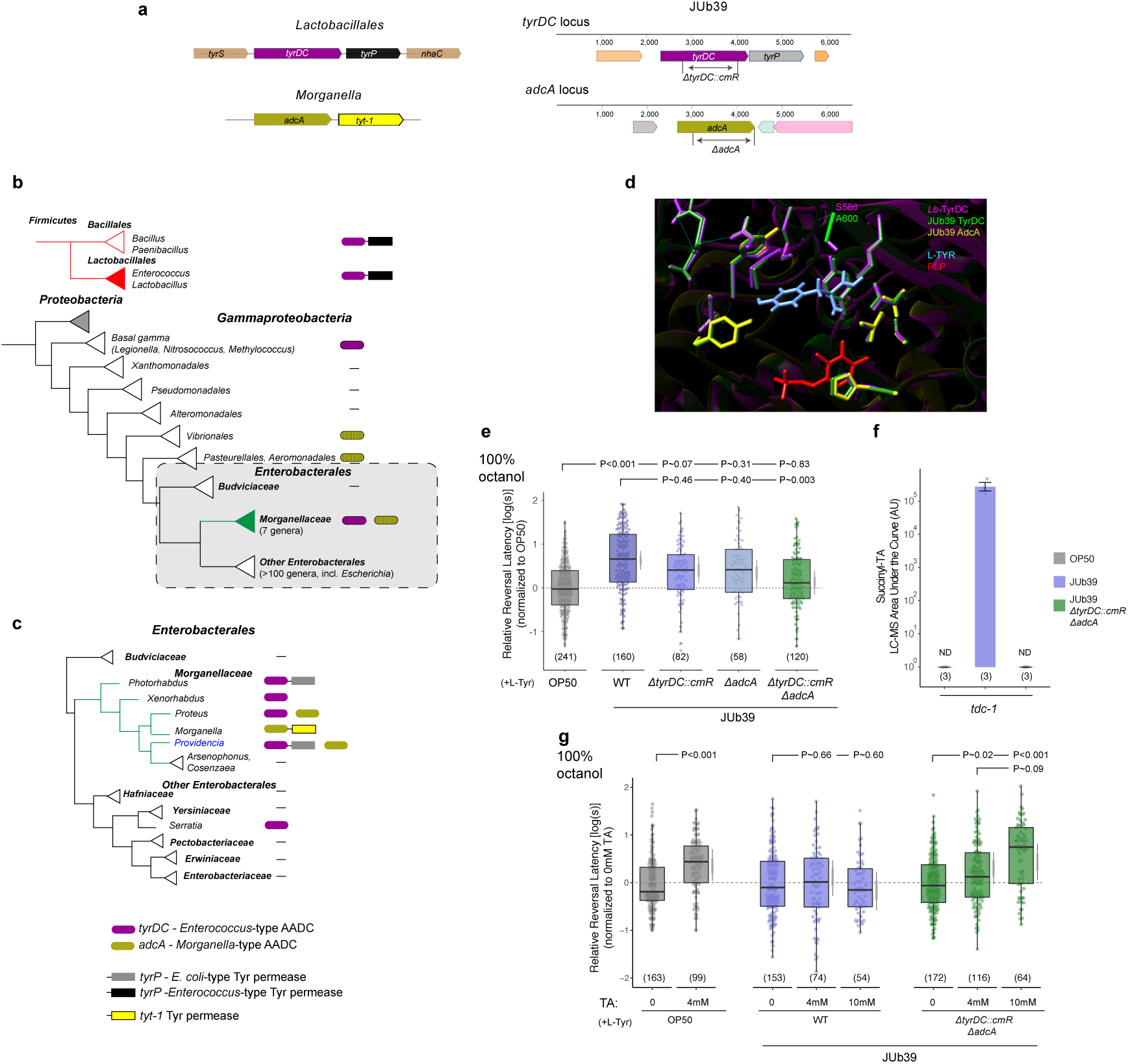
Two amino acid decarboxylase enzymes encoded by the *Providencia* genome act redundantly to modulate octanol avoidance. **a)** Cartoons depicting the *tyrDC* locus (top) in *Lactobacillales* (left) and JUb_39_ (right) and the *adcA* locus (bottom) in *Morganella* (left) and JUb_39_ (right). **b)** Presence of *tyrDC* and *adcA* among complete genomes in *Gammaproteobacteria*. Linked boxes indicate organization in an operon. Hatched shading indicates variable presence among genera. Colored triangles indicate taxa of interest. **c)** Presence of *tyrDC, adcA, E.coli*-type *tyrP* and *Morganella*-type *tyt-1* at the family and genus level among *Enterobacteriales*. Linked boxes indicate organization in an operon. **d)** Homology-based model of the TyrDC catalytic domain in *Providencia* based on the *Lb*-TyrDC crystal structure^34^ using SWISS-MODEL (https://swissmodel.expasy.org). Residues in magenta, green and yellow are from *Lb*-TyrDC, JUb_39_-TyrDC, and JUb_39_-AdcA, respectively. PLP is depicted in red and L-Tyr (manually docked for illustration) is indicated in light blue. Position of A600/S586^34^in JUb_39_ TyrDC and *Lb*-TyrDC are indicated. **e, g)** Reversal response latency of animals of wild-type *C. elegans* grown on the indicated bacterial genotypes in either control conditions of NGM + 0.5% L-Tyr **(e)** or supplemented with the indicated concentrations of TA **(g)** to 100% octanol using SOS assays. Each dot is the response time of a single worm. Y-axis is log_10_-scaled for these log-normal distributed data, and normalized to the indicated control group for each experimental day, indicated by the gray horizontal line. Numbers in parentheses indicate the number of worms tested in assays over at least 3 independent days. Boxplot indicates median and quartiles, whiskers indicate the data range, excluding outliers. Gray thin and thick vertical bars at right indicate Bayesian 95% and 66% credible intervals for the difference of means, respectively. *P*-values between indicated conditions are from a LMM with Tukey-type multivariate-t adjustment. **f)** Quantification of succinyl-TA in *tdc-1* mutant animals grown on the indicated bacterial strains. OP_50_ and JUb_39_ data are repeated from Fig. 2d. Data are averaged from three independent replicates each. ND, not detected.

*Providencia* TyrDC is highly homologous to the *Lb* enzyme, which has been well characterized with respect to substrate specificity^34^. Protein modeling using the crystal structure of *Lb*-TyrDC^34^ as a guide (see Methods) indicated that JUb_39_ TyrDC shares most known catalytic sites with *Lb*-TyrDC (Fig. 3d). Interestingly, JUb_39_ TyrDC contains a substitution at A600 (S586 in *Lb*-TyrDC; Fig.3d), a variant demonstrated to enhance specific catalytic activity of *Lb*-TyrDC for tyrosine^34^. We infer that JUb_39_ TyrDC likely generates TA from tyrosine.

*Morganella* strains (*Morganellaceae* family) have been reported to produce TA under certain conditions^28^, despite having no discernible tyrDC orthologs (Fig. 3c, Fig. S4a-b, Table S1). Instead in *Morganella*, we identified an AADC-encoding gene (hereafter *adcA*) with ∼29% and 27% sequence identity to *Enterococcus* TyrDC and human GAD67, respectively, in an operon upstream of a gene encoding a TYT-1 family tyrosine permease (Fig. 3a, Fig. 3c). An *adcA* homolog is also present in *Providencia* genomes including in JUb_39_ but is not adjacent to a tyrosine transporter (Fig. 3a, Fig. S4a-b, Table S1). We conclude that *Providencia* encodes at least two AADCs with the potential to generate TA, and the phylogenetic incongruence suggests that both tyrDC and *adcA* genes may have either been lost or acquired in the *Morganellaceae* family via horizontal gene transfer.

To test whether one or both JUb_39_ AADCs are necessary for octanol modulation, we engineered deletions in JUb_39_ *tyrDC* and *adcA* (Δ*tyrDC::cmR* and Δ*adcA*, respectively; Fig. 3a). While cultivation on each deletion-containing bacterial strain alone weakly decreased octanol modulation, growth on the JUb_39_ Δ*tyrDC::cmR* Δ*adcA* double knockout bacteria abolished octanol modulation (Fig. 3e). We confirmed that JUb_39_ Δ*tyrDC::cmR* Δ*adcA* colonizes the *C. elegans* gut (Fig. S5), and further showed that *tdc-1* animals grown on JUb_39_ Δ*tyrDC::cmR* Δ*adcA* do not produce succinyl-TA (Fig. 3f). Octanol modulation was restored in wild-type *C. elegans* grown on JUb_39_ Δ*tyrDC::cmR* Δ*adcA* strains upon supplementation with TA (Fig. 3g). Moreover, while exogenous TA did not further increase octanol avoidance in wild-type JUb_39_-grown animals, TA supplementation was sufficient to induce octanol modulation in OP_50_-grown animals (Fig. 3g). Together, these results indicate that TA produced by multiple AADC enzymes in *Providencia* is both necessary and sufficient to modulate octanol avoidance by wild-type *C. elegans*.

We next identified the molecular targets of *Providencia*-mediated octanol modulation in the host. As bacterially-produced TA is likely converted to OA via the host TBH-1 enzyme^23^ to mediate octanol modulation, we focused primarily on host OA receptors. The bilateral ASH nociceptive neurons located in the head amphid organs of *C. elegans* have been implicated in sensing octanol^9,19,26^. These neurons express multiple TA and OA receptors, a subset of which is required for octanol modulation by these monoamines^18,35^. Among ASH-expressed OA receptors, mutations in *octr-1*, but not *ser-3*, abolished JUb_39_-mediated octanol modulation, without altering the extent of gut colonization (Fig. 4a, Fig. S5). We also observed an effect on octanol modulation in *tyra-2* TA receptor mutants, primarily due to decreased octanol avoidance upon growth on OP_50_ (4.3 ± 0.25s for *tyra-2* vs. 2.9 ± 0.13s for WT). TYRA-2 has recently been shown to mediate responses to an OA-linked pheromone^36^, although the reason for the observed effect in OP_50_-grown animals is currently unclear. Expression of *octr-1* cDNA in the ASH/ASI sensory neurons restored octanol modulation (Fig. 4a). *octr-1* mutants also lacked octanol modulation when grown on JUb_39_ Δ*tyrDC::cmR* Δ*adcA* supplemented with TA (Fig. 4b). We conclude that host OA produced upon JUb_39_ colonization acts via OCTR-1 in ASH/ASI to modulate octanol avoidance.

**Fig. 4.**
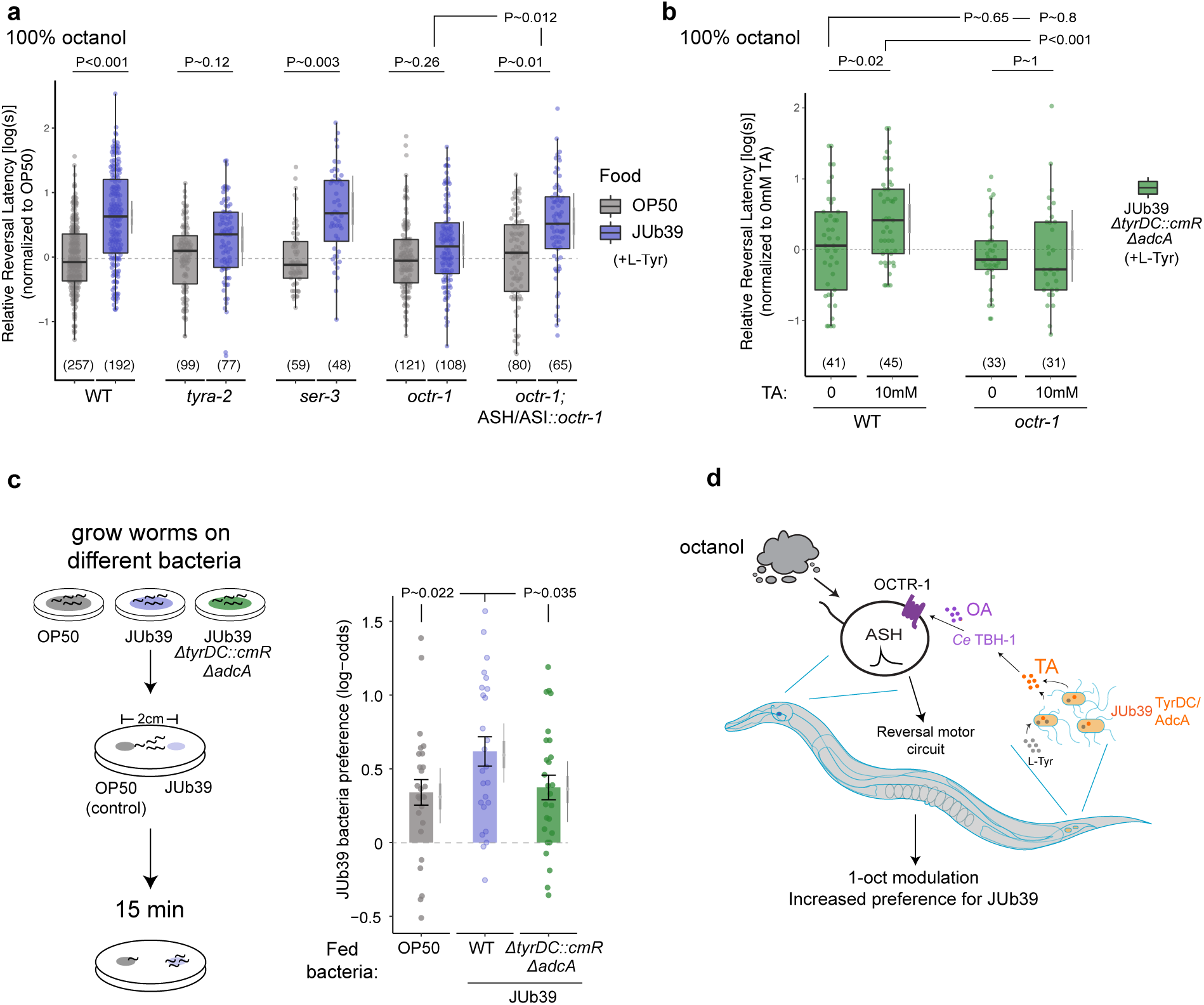
Modulation of octanol avoidance by *Providencia* requires the OCTR-1 OA receptor in the ASH/ASI sensory neurons. **a-b)** Reversal response latency of animals of the indicated genotypes grown on the shown bacteria in control conditions of NGM + 0.5% L-Tyr **(a)** or supplemented with TA + 0.5% L-Tyr **(b)** to 100% octanol using SOS assays. Each dot is the response time of a single worm. Y-axis is log_10_-scaled for these log-normal distributed data, and normalized to the indicated control group for each experimental day. Numbers in parentheses indicate the number of worms tested in assays over at least 3 independent days. Boxplot indicates median and quartiles, whiskers indicate the data range, excluding outliers. Gray thin and thick vertical bars at right indicate Bayesian 95% and 66% credible intervals for the difference of means, respectively. *P*-values between indicated conditions are from a LMM with Tukey-type multivariate-t adjustment. **c)** (Left) Cartoon depicting assay setup of the short-range bacterial choice assay. (Right) Preference index of animals grown on the indicated bacteria for the test bacteria JUb_39_. Each dot represents one assay of at least 10 animals; assays were performed over at least 4 independent days. Y-axis is on log-odds (logit) scale. Errors are SEM. Gray thin and thick vertical bars at right indicate Bayesian 95% and 66% credible intervals, respectively. *P*-values represent difference of means relative to JUb_39_-grown animals from a GLMM with Dunnett-type multivariate-t adjustment. **d)** Cartoon of working model. JUb_39_ colonizes the *C. elegans* intestine and produces TA via the TyrDC and AdcA enzymes. TA is converted to OA by *C. elegans* TBH-1 and acts via the ASH neuron-expressed OCTR-1 OA receptor to modulate octanol avoidance and food choice.

Next we investigated the biological relevance of the JUb_39_-directed decrease in octanol aversion by *C. elegans*. While many gram-negative enteric bacteria produce long-chain alcohols including octanol^37^, whether *Providencia* produces this chemical is unknown. However, JUb_39_ and other *Providencia* strains can produce the branched alcohol isoamyl alcohol (IAA), which is aversive to *C. elegans* when concentrated^38,39^. Similar to octanol, avoidance of high IAA concentrations is also mediated by the ASH sensory neurons^39^. We hypothesized that reduced avoidance of JUb_39_-produced aversive alcohols or other odorants may preferentially bias JUb_39_-grown *C. elegans* to select these bacteria in food choice assays. Indeed, animals grown on JUb_39_ more strongly preferred JUb_39_ compared to OP_50_-grown worms, which showed a slight preference for JUb_39_ in a short-range food choice assay (Fig. 4c). The bias towards JUb_39_ was eliminated in animals grown on JUb_39_ Δ*tyrDC*::cmR Δ*adcA*, suggesting that bacterial TA production is necessary for this food preference (Fig 4b). Together, these results imply that TA produced by JUb_39_ reduces ASH/ASI-mediated avoidance of JUb_39_-produced aversive cues such as concentrated alcohols, to allow preferential selection of these bacteria.

Our observations support a model in which the neurotransmitter TA produced by intestinal *Providencia* bacteria modulates aversive responses of *C. elegans* to the enteric bacteria-produced volatile metabolite octanol, likely via subverting host-dependent TA production. Bacterially-produced TA is converted to OA by *C. elegans* TBH-1; OA subsequently acts on the ASH/ASI neurons via the OCTR-1 OA receptor to decrease aversion of octanol. Bacterially-derived TA also increases the preference of *C. elegans* for *Providencia* in food choice assays (Fig. 4d). We speculate that the preference for *Providencia* upon colonization of *C. elegans* by these bacteria promotes increased consumption leading to stable association^4,40^ and bacterial dispersal. As *Providencia* is a rich food source for *C. elegans*^6^, this association may be mutually beneficial. Our results describe a pathway by which neurotransmitters produced by commensal bacterial direct host behavioral decisions by supplementing or compensating for the activity of key host biosynthetic enzymes, thereby altering fitness of both host and microbe.

## Methods

### Strains

#### C. elegans

All *C. elegans* strains were maintained on nematode growth medium (NGM) at 20°C. *sra-6p::octr-1* plasmid (pMOD100) was injected at 10 ng/*µ*l together with the *unc-122p::gfp* coinjection marker at 30 ng/*µ*l to generate transgenic strains. Two independent lines were examined for *octr-1* rescue experiments.

#### Bacteria

For all experiments, bacterial strains were streaked from glycerol stocks prior to use and grown to saturation in LB media at 37°C. For conditioned media, bacteria were grown to saturation in NGM media overnight at 37°C, then cleared by centrifugation at 14,000g for 3 minutes. Prior to use, conditioned media or NGM was supplemented with 5x concentrated OP_50_ from a saturated LB culture to prevent starvation. To expose animals to bacterial odors, worms were grown on seeded NGM plates whose lids were replaced with NGM plates containing the test bacteria; these were sealed with parafilm. For L-Tyr and TA supplementation experiments, 0.5% L-Tyr (Sigma T3754) or 4mM or 10mM TA (Sigma T2879) were added to the NGM media and agar prior to pouring plates. Plasmids were transformed into JUb_39_ and OP_50_ via electroporation. Deletions in JUb_39_ were induced using homologous recombination with the temperature-sensitive pSC101 replicon at 42°C, and *sacB*-sucrose counter-selection at 30°C, in the absence of NaCl as described^41^, with the exception that bacteria were incubated for 1 hour at room temperature in the presence of 10mM arabinose for lambda Red induction prior to selection at 42°C. Deletions were confirmed by sucrose resistance and kanamycin sensitivity, followed by PCR and sequencing of deleted intervals.

### Molecular biology

The *octr-1* cDNA was a gift from Dr. Richard Komuniecki. The cDNA was amplified by PCR and cloned using Gibson homology cloning. The 3.8kb *sra-6* promoter sequence was cloned from genomic DNA. Vector maps are available on Github (https://github.com/SenguptaLab/ProvidenciaChemo.git). For introduction of deletions via homologous recombination in JUb_39_, pKD46-derivative plasmids containing a lambda Red cassette and deletion homology arms for JUb_39_ *tyrDC* and *adcA* were constructed (denoted pMOD102 and pMOD107, respectively). Briefly, the *cas9* coding region and sgRNA regions of pDK46-derivative pCAS^42^ were deleted and replaced with the *sacB* sequence from pCM433^43^ via PCR and Gibson homology cloning. For pMOD102, 5’ and 3’ homology arms were approximately 400bp each flanking a 1233bp deletion of the *tyrDC* coding sequence which was replaced with a chloramphenicol resistance cassette. For pMOD107, 5’ and 3’ homology arms were 701 and 422bp, respectively, flanking a 1398bp deletion of the *adcA* CDS. For expression of mCherry in OP_50_ and JUb_39_, a pUCP20T-mCherry plasmid^44^ was modified to replace *bla*(ampR) with *aph*(kanR).

### Microscopy

All fluorescence microscopy was performed using animals anesthetized with 100 mM levamisole (Sigma Aldrich). Animals were imaged on 2% agarose pads using an upright Zeiss Axio Imager with a 63X oil immersion objective.

#### Quantification of intestinal bacterial cell numbers

All rod-shaped punctae in the intestines of young adult worms of approximately 1-2*µ*m were included in the quantification. Each animal was recorded in one of three categories containing 0, <10, or >10 cells per animal. Exact numbers in animals bearing over 10 cells were not recorded, but rarely exceeded approximately 100 cells.

#### Fluorescence intensity measurements

All images were collected in z-stacks of 0.5 *µ*m through the heads of young adult worms. Quantification was performed using ImageJ (NIH). Fluorescence was quantified by identifying the focal plane in which the cell soma was visible, followed by manually drawing an ROI around the soma. Mean pixel intensity was recorded for each neuron pair per animal and the average of fluorescence in each animal is shown.

### Behavioral assays

#### Long-range chemotaxis

Long-range chemotaxis assays were performed essentially as described^7,45^. Worms were cultured for 1 generation with the relevant bacteria prior to the assay. Assays were performed using 10cm square NGM plates. The number of worms in two horizontal rows adjacent to the odor and ethanol spots were quantified.

#### SOS assays

Smell-on-a-stick (SOS) assays in response to 1-octanol or 2-nonanone were performed as described^9,26^. NGM plates were pre-dried for 1 hour prior to assays. Age-matched young adult animals were picked from food to a clean transfer plate and allowed to briefly crawl away from food for approximately 1 min. Animals were then transferred to another clean NGM plate for 15 minutes prior to assaying responses to 100% octanol (Sigma O4500) and 100% 2-nonanone (Sigma 108731), or 20 minutes for 30% octanol assays. 30% octanol was prepared immediately before the assay by dilution in 200-proof ethanol (Acros Organics 61509-0010).

#### Short-range bacterial choice assay

Animals were raised and prepared identically to those used in long-range chemotaxis assays, with the exception that the final wash with water was omitted. NGM plates containing 2 15*µ*L spots of overnight-grown bacterial food concentrated to OD600 ∼ 10 placed 2cm apart were allowed to dry, then incubated with a closed lid for 5 hrs at room temperature. Approximately 30 animals were placed between the two spots, and excess liquid was removed. Animals were allowed to navigate for 15 minutes following which 2*µ*L of sodium azide was applied to each spot to anesthetize worms. Very little lawn-leaving behavior was observed during this short time period. Adult animals on the control spot and test spot were counted.

#### Osmotic avoidance assay

Animals off the bacterial food on the cultivation plate were picked using a 10% methyl cellulose polymer solution and placed in the center of an NGM plate with a ring of 8M glycerol containing bromophenol blue (Sigma B0126). The number of worms inside and outside of the ring were counted after 10 mins.

### Bacteria genome sequencing

Sequencing was performed by the Broad Technology Labs at the Broad Institute. Resulting PacBio reads for JUb_39_ and PYb_007_ were assembled using Canu^46^ v1.8 (https://github.com/marbl/canu.git). Assemblies were trimmed, oriented and circularized using Circlator^47^ v1.5.5 (https://sanger-pathogens.github.io/circlator/).

### Phylogenetic analysis of group II pyridoxal-dependent decarboxylase genes

JUb_39_ TyrDC and AdcA were initially identified as the only significant hits via a tblastn search of the draft JUb_39_ genome assembly using *Enterococcus faecalis* TyrDC as a query sequence. An initial BLASTP screen of the nr sequence database restricted to bacteria was performed using the *P. alcalifaciens* JUb_39_ TyrDC and AdcA coding regions. Searches were performed hierarchically, limited initially to *Enterobacteriaceae*, followed by *Enterobacterales, Gammaproteobacteria, Proteobacteria* and finally all *Eubacteria*. With the exception of members of *Morganellaceae* (*Providencia, Proteus, Morganella, Xenorhabdus, Photorhabdus, Arsenophonus* and *Moellerella*), only two protein sequences per genus were retained for subsequent phylogenetic analysis. Representative group II decarboxylase enzymes with known substrate specificity from *Eukaryota* and *Archaea* as well as glutamate decarboxylase (GadA/B) and histidine decarboxylase sequences were also included.

Multiple sequence alignments were produced using the Phylomizer workflow (https://github.com/Gabaldonlab/phylomizer), which used the MUSCLE^48^ v3.8.31 (http://www.drive5.com/muscle), MAFFT^49^ v7.407 (https://mafft.cbrc.jp/alignment/software) and Kalign^50^ v2.04 (http://msa.sbc.su.se/cgi-bin/msa.cgi) multiple sequence aligners; these were trimmed to produce a consensus alignment using trimAL^51^ v1.4rev15 (https://github.com/scapella/trimal). An initial phylogenetic tree was produced using PhyML^52^ v3.3.20180621 (http://www.atgc-montpellier.fr/phyml/) using the NNI algorithm with an LG substitution model. This tree showed three major, well-supported clusters containing: (1) *Enterococcus* and *Providencia* TyrDCs - denoted “*Enterococcus*-type TDC”, (2) Eukaryotic AADCs denoted “Eukaryotic-type AADC”, and (3) *Morganella* AdcA and *Providencia* AdcA.

Based on this initial tree, a second tblastn search was used to determine the presence or absence of homologous genes among complete *Gammaproteobacteria* genomes. *Enterococcus faecalis* TyrDC and *C. elegans* TDC-1 were used as tblastn search query sequences. Hierarchical search was performed as described above, limited to an e-value cutoff of 10-5. A maximum of 2 highly similar sequences were retained per genus for phylogenetic analysis as listed in Table S1.

A final phylogenetic tree was constructed using the amino acid sequences derived from these tblastn queries. These were assembled into a consensus alignment using the Phylomizer workflow as described above. ProtTest^53^ (https://github.com/ddarriba/prottest3) was used to identify the optimal model for likelihood estimation, using Aikake Information Criterion (AIC) values for selection. The model selected and subject to PhyML analysis was an LG model with discrete gamma distribution, an estimated proportion of invariant sites (+I), empirical frequencies of amino acids (+F), estimated gamma shape parameter (+G) for rate variation among sites with the default 4 substitution rate categories, and the subtree pruning and regrafting (SPR) algorithm. 100 bootstrap pseudoreplicates were analyzed. Representatives from the resulting phylogeny were used to categorize and compile the cladogram in Fig. 3b-c. Adjacent genomic sequences, up to 3 CDS 5’ or 3’, were examined for genes encoding amino-acid permeases or transporters in an apparent operon as defined by close proximity and same orientation with respect to each tblastn hit (Table S1).

### Molecular modeling

The putative amino acid sequence for JUb_39_ TyrDC was used to model active site residues using the *Lb*-TyrDC crystal structure in complex with PLP (5hsj.1^34^) as a template guide using SWISS-MODEL (https://swissmodel.expasy.org). This resulted in a Qmean Z-score of 0.33, indicative of good agreement between structures. This process was also attempted with AdcA, and modeling was performed with the top 6 available structures based on sequence homology. The maximum QMean of AdcA was found with *Lb*-TyrDC, but with a value −5.71, indicative of low quality. Resulting models were visualized using Chimera^54^ v1.13.1 (https://www.cgl.ucsf.edu/chimera/). For Fig. 3d, L-Tyrosine was manually docked according to the reported docking position^34^ for illustrative purposes only.

### Statistical analyses

All statistical analyses were performed in R (https://www.R-project.org/) and RStudio (http://www.rstudio.com). For modulation index and relative latency figures, data were were normalized to the relevant control group mean value for each experimental day on the log scale via subtraction. Outliers in boxplots were defined as greater than 1.5*interquartile range, but were included for analysis. All statistical analyses were performed on raw, non-normalized data. To avoid inflated *P*-values and account for non-independence of observations, we employed mixed-effects regression analysis in lieu of simple ANOVA and t-tests. For behavioral assays, frequentist statistical comparisons were performed using a binomial generalized linear mixed-effects model (GLMM) with a logit link function for chemotaxis and food choice assays, while a linear mixed-effects model (LMM) on log_10_-transformed data was used to analyze SOS assays using the ‘lme4’ package. In all cases, a random intercept term for assay plate was used to account for non-independence of animals on each assay plate and random intercept for date was used to account for day-to-day variability. In the presence of interactions, for example the effects of bacterial strains across different odorants in Fig. 1a, a random slope term per date was also used when appropriate. Estimated *P*-values for pairwise comparison of fixed effects were determined using Kenward-Roger approximated degrees of freedom as implemented in the ‘emmeans’ and ‘pbkrtest’ packages. In nearly all cases, inclusion of random effects model terms resulted in conservative *P*-value estimates compared to a simple ANOVA. In the event of singular model fit, any random slope term, followed by random date effect terms were removed to allow convergence. For Wald statistics of model terms, packages ‘lmerTest’ or ‘car’ were used.

Additionally, for each dataset, a maximal Bayesian model was fit using the ‘rstanarm’ and ‘rstan’ packages. Data presented are posterior credible intervals for fixed effect levels derived from posterior fitted values of the MCMC chains as implemented by the ‘emmeans’, ‘coda’, ‘bayesplot’ and ‘tidybayes’ packages. Post-hoc corrections for multiple comparisons and type-I error were implemented using the ‘emmeans’ package. For comparison of intestinal bacterial cell numbers, an ordinal logistic regression was performed using the ‘MASS’ package and ‘polr’ function. Categories of cell numbers were considered ordered factors of ‘none’, ‘some’ or ‘many’ cells.

### Sample preparation for HPLC-MS

Approximately 10,000 mixed-staged worms in 1.5mL microfuge tubes were lyophilized for 18-24 hrs using a VirTis BenchTop 4K Freeze Dryer. After the addition of two stainless steel grinding balls and 1mL of 80% methanol, samples were sonicated for 5 min (2 sec on/off pulse cycle at 90 A) using a Qsonica Q700 Ultrasonic Processor with a water bath cup horn adaptor (Model 431C2). Following sonication, microfuge tubes were centrifuged at 10,000 RCF for 5 min in an Eppendorf 5417R centrifuge. 800*µ*L of the resulting supernatant was transferred to a clean 4mL glass vial, and 800*µ*L of fresh methanol added to the sample. The sample was sonicated and centrifuged as described, and the resulting supernatant was transferred to the same receiver vial and concentrated to dryness in an SC250EXP Speedvac Concentrator coupled to an RVT5105 Refrigerated Vapor Trap (Thermo Scientific). The resulting powder was suspended in 120*µ*L of 100% methanol, followed by vigorous vortex and brief sonication. This solution was transferred to a clean microfuge tube and subjected to centrifugation at 20,000 RCF for 10 min in an Eppendorf 5417R centrifuge to remove precipitate. The resulting supernatant was transferred to an HPLC vial and analyzed by HPLC-MS.

### HPLC-MS analyses

Reversed-phase chromatography was performed using a Vanquish LC system controlled by Chromeoleon Software (ThermoFisher Scientific) and coupled to an Orbitrap Q-Exactive High Field mass spectrometer controlled by Xcalibur software (ThermoFisher Scientific). Methanolic extracts prepared as described above were separated on an Agilent Zorbax Eclipse XDB-C18 column (150 mm x 2.1 mm, particle size 1.8 *µ*m) maintained at 40°C with a flow rate of 0.5mL/min. Solvent A: 0.1% formic acid in water; solvent B: 0.1% formic acid in acetonitrile (ACN). A/B gradient started at 5% B for 3 min after injection and increased linearly to 98% B at 20 min, holding at 98% B until 25 min, followed by a 0.1 min gradient to 5% B, held until 28 min. Mass Spectrometer parameters: spray voltage 3.0 kV, capillary temperature 380°C, probe heater temperature 400°C; sheath, auxiliary, and sweep gas 60, 20, and 1 AU, respectively. S-Lens RF level: 50, resolution 240,000 at m/z 200, AGC target 3E6, maximum injection time (IT) 500 msec. Each sample was analyzed in negative and positive electrospray ionization modes with m/z range 150–800. Parameters for MS/MS (dd-MS2): MS1 resolution: 60,000, AGC Target: 3E6, max IT 100 msec. MS2 resolution: 30,000, AGC Target: 1E5, max IT 50 msec. Isolation window 1.0 m/z, stepped normalized collision energy (NCE) 10, 30; dynamic exclusion: 5 sec. Top 10 masses selected for MS/MS per scan. LC-MS data were analyzed via manual integration in Excalibur (Thermo Fisher Scientific).

## Data availability

All statistical analysis code and raw data necessary to reproduce these analyses are available [https://github.com/SenguptaLab/ProvidenciaChemo.git]. Draft genome assemblies will be deposited in Genbank.

## Acknowledgements

We thank Richard Komuniecki for the *octr-1* cDNA, multiple members of the *C. elegans* community for bacterial strains (listed in Table S2), and the *Caenorhabditis* Genetics Center for *C. elegans* strains. We are grateful to the Broad Technology Labs for bacterial genome sequencing. We thank Sue Lovett, Laura Laranjo and the Sengupta lab for advice. We acknowledge the Sengupta lab and Cori Bargmann for comments on the manuscript. This work was partly supported by the NIH (T32 NS007292 and F32 DC013711 – M.O’D, R01 GM088290 and R35 GM131877 – F.C.S., and R35 GM122463 and R21 NS101702 – P.S.), and the NSF (IOS 1655118 – P.S.).

