## Supplemental Figures for "Modulation of sensory behavior and food choice by an enteric bacteria-produced neurotransmitter"

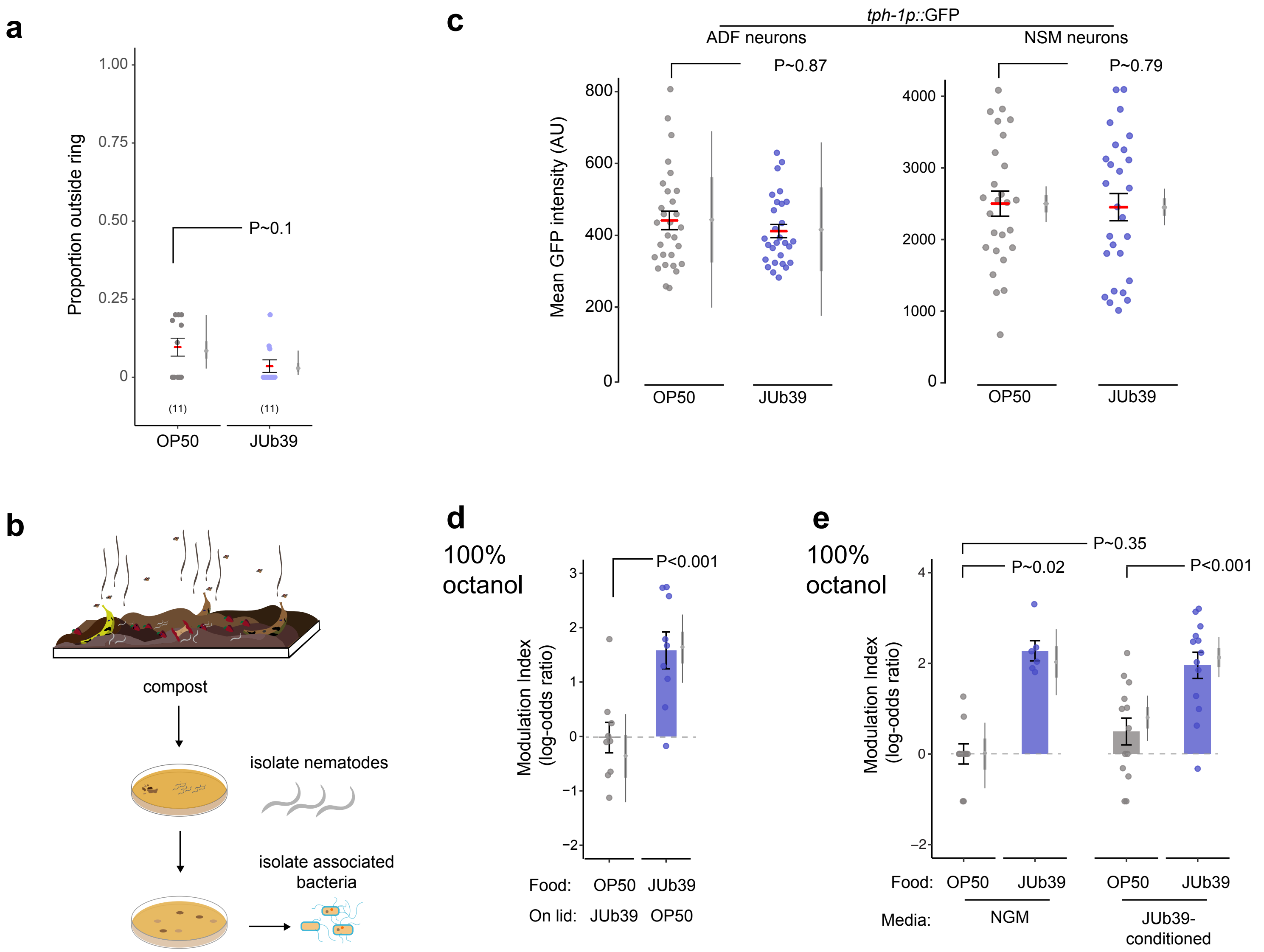


**Fig. S1. Octanol modulation by *Providencia* requires ingestion of bacteria and is not mediated by nutritive cues.**
**a)** Osmotic ring avoidance assays. Each dot represents one assay of 10 animals. Numbers in parentheses indicate the number of assays over at least 3 independent days. Y-axis is proportion of animals leaving an osmotic ring barrier of 8M glycerol after 10 minutes. $P$-value represents difference of means relative to JUb39-grown animals from a GLMM. Errors are SEM. Gray thin and thick vertical bars at right indicate Bayesian 95% and 66% credible intervals, respectively.
**b)** Isolation of nematode-associated bacteria. Nematodes were isolated from residential compost in Massachusetts. Worms were allowed to crawl onto NGM plates from which they were picked to clean plates. Resulting bacterial colonies were isolated, grown on LB media and characterized via 16S rRNA sequencing.
**c)** Expression of a *tph-1p::gfp* fluorescent reporter in indicated head neurons of young adult animals grown on either OP50 or JUb39. Each dot is the mean fluorescence of the soma of one neuron. Horizontal bar is mean; errors are SEM. Gray thin and thick vertical bars at right indicate Bayesian 95% and 66% credible intervals, respectively. $P$-values are from two-way ANOVA.
**d-e)** Modulation index of worms grown on the indicated bacterial strains, under the shown conditions. Animals were exposed to the indicated bacteria on the plate lid **(d)** for one generation, or to NGM control or bacteria-conditioned NGM **(e)** for 2 hours prior to the assay. Each dot represents results from one chemotaxis assay with approximately 100 animals each. Values are shown on a log-odds (logit) scale and are normalized to the values of wild-type animals grown on OP50 for each day, indicated with a gray dashed line. Positive numbers indicate reduced avoidance of octanol. Errors are SEM. Gray thin and thick vertical bars at right indicate Bayesian 95% and 66% credible intervals, respectively. $P$-values between the indicated conditions are post-hoc comparisons from a GLMM, with Tukey-type multivariate-t adjustment for **e**.

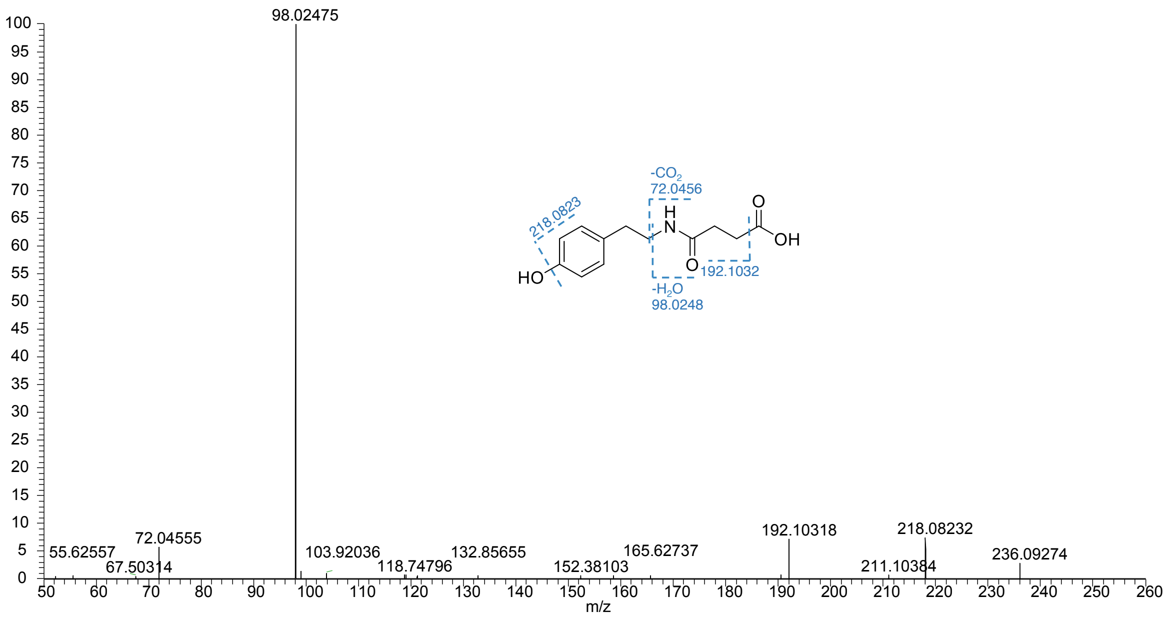


**Fig. S2. Structure of succinyl-TA.**
High-resolution MS/MS spectrum of succinyl-TA in negative ionization mode. Structures of major MS/MS fragmentations are denoted by dashed lines.

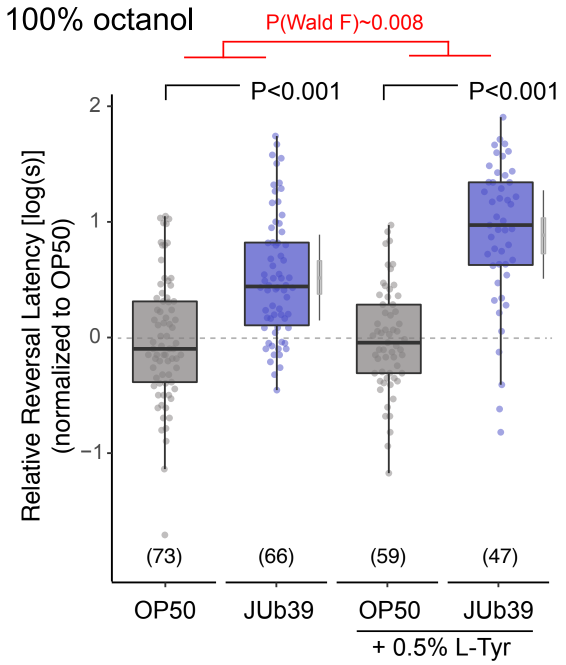


**Fig. S3. L-Tyr supplementation enhances octanol modulation.**
Reversal response times of animals of the indicated genotypes grown on the indicated bacteria in control conditions or supplemented with 0.5% L-Tyr to 100% octanol using SOS assays. Each dot is the response time of a single worm. Y-axis is $\log_{10}$-scaled for these log-normal distributed data, and normalized to the indicated control group for each experimental day. Numbers in parentheses indicate the number of worms tested in assays over at least 3 independent days. Boxplot indicates median and quartiles, whiskers indicate the data range, excluding outliers. Gray thin and thick vertical bars at right indicate Bayesian 95% and 66% credible intervals for the difference of means, respectively. *P*-values indicating comparisons of means relative to the OP50 control for each conditions are from a LMM with Tukey-type multivariate-t adjustment. *P*-value in red indicates Wald F-statistic for the effect of L-Tyr supplementation on the magnitude of the JUb39 effect.

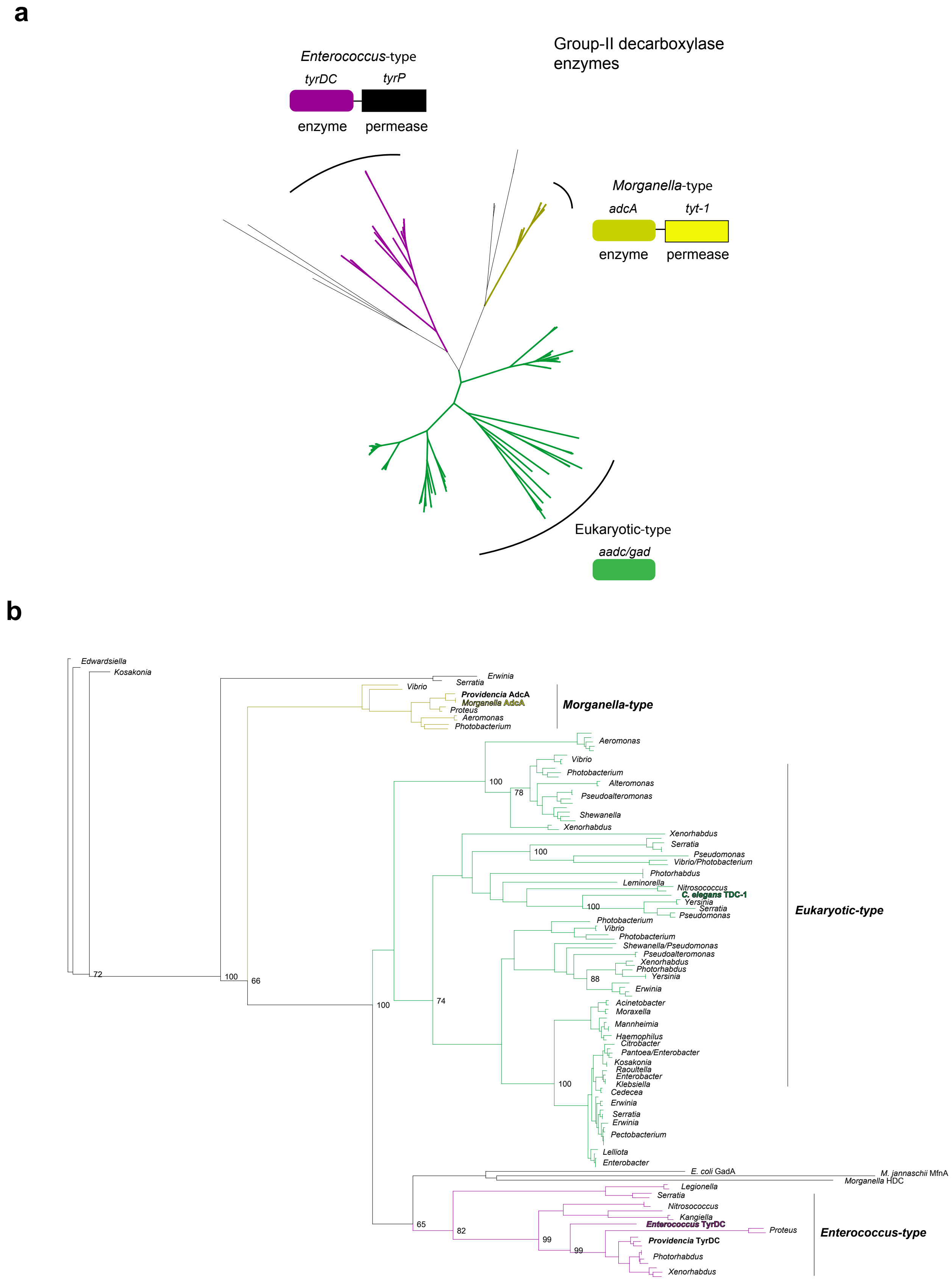


**Fig. S4. Phylogenetic analysis of group II decarboxylase genes in *Gammaproteobacteria*.**

**a)** Neighbor-joining unrooted tree based on sequences identified via a BLAST search using *Enterococcus faecalis* TyrDC and *C. elegans* TDC-1. Initial tree indicates 3 major groups. Representative enzymes and operon structures for each group are indicated by colored boxes.

**b)** Bootstrapped maximum likelihood phylogeny using PhyML and Phylomizer pipeline. Maximum of two highly similar sequences per genus were included after each BLAST search. Genera are indicated to the right. Numbers on branch-points matching this tree out of 100 bootstrap replicates are indicated at values >60. Group representatives from **(a)** are indicated in corresponding colors. *Providencia* and *C. elegans* sequences discussed in this work are indicated in bold. Accession numbers and BLAST metrics are listed in Extended Data Table 1.

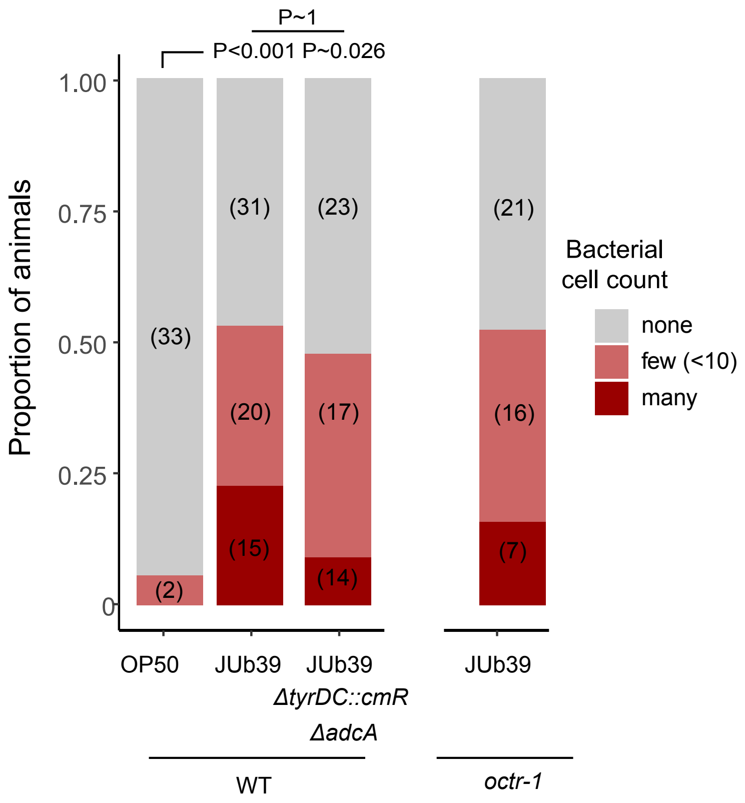


**Fig. S5. Mutations in *Providencia* TDC-encoding genes or *C. elegans octr-1* do not alter intestinal *Providencia* numbers.**
Presence of mCherry-expressing bacteria in the posterior intestines of young adult wild-type or *octr-1* mutant animals. Bars show proportion of animals with the indicated distribution of bacterial cells present in animals grown on shown bacteria. Numbers in parentheses indicate the number of animals. *P*-value is derived from an ordinal regression.
