## Supplemental Table 2 for "Modulation of sensory behavior and food choice by an enteric bacteria-produced neurotransmitter"

**Extended Data Table 2.** Strains used in this work.

| ***C. elegans* strain** | **Genotype** | **Source and/or**  **parent strains^a,b^** |
| --- | --- | --- |
| N2 | N2 (Bristol) WT | CGC |
| RB993 | *tdc-1(ok914)* II | CGC |
| RB1161 | *tbh-1(ok1196)* X | CGC |
| MT15620 | *cat-2(n4547)* II | CGC |
| MT15434 | *tph-1(mg280)* II | CGC |
| PY9349 | *tdc-1(ok914)* II; *tbh-1(ok1196)* X | RB993 and RB1161 |
| PY9350 | *tyra-2(tm1815)* X  (backcrossed 2X) | NBRP |
| CX12800 | *ser-3(ad1774)* I | CGC |
| CX13079 | *octr-1(ok371)* X | CGC |
| PY9351 | *octr-1(ok371)* X; *oyEx630*[*sra-6*p*::octr-1*, *unc-122*p*::mcherry*] line 1 | pMOD100 injected into CX13079 |
| PY9352 | *octr-1(ok371)* X; *oyEx631*[*sra-6*p*::octr-1*, *unc-122*p*::mcherry*] line 2 | pMOD100 injected into CX13079 |
| GR1333 | *yzIs71*[*tph-1*p*::gfp, rol-6(su10060*] | CGC |
| **Bacterial Strain** | **Species/Genotype** | **Source and/or**  **parent strains** |
| OP50 | *E. coli /* WT | Laboratory strain, CGC |
| JUb39 | *Providencia alcalifaciens* / WT | *C. elegans* in rotting apple (isolated by Marie-Anne Félix)^1,2^ |
| DA1877 | *Comomonas sp.* / WT | Soil (isolated by Boris Shtonda)^3^ |
| DA1878 | *Pseudomonas sp.* / WT | Soil (isolated by Boris Shtonda)^3^ |
| DA1880 | *Bacillus megaterium* / WT | Soil (isolated by Boris Shtonda)^3^ |
| PYb007 | *Providencia rettgeri* / WT | Nematodes isolated from residential compost |
| PA103 | *Pseudomonas aeruginosa* | Laboratory strain, gift from Yun Zhang^4^ |
| PAK | *P. aeruginosa* | Laboratory strain, gift from Yun Zhang^4^ |
| PYb110 | *P. alcalifaciens ΔtyrDC*::cmR | Homologous recombination in JUb39 |
| PYb111 | *P. alcalifaciens ΔadcA* | Homologous recombination in JUb39 |
| PYb112 | *P. alcalifaciens ΔtyrDC*::cmR *ΔadcA* | Homologous recombination in PYb110 |

^a^CGC – *Caenorhabditis* Genetics Center; ^b^NBRP – National BioResource Project

1.	Samuel, B. S., Rowedder, H., Braendle, C., Félix, M.-A. & Ruvkun, G. *Caenorhabditis elegans* responses to bacteria from its natural habitats. *Proc. Natl. Acad. Sci.* **113,** E3941–9 (2016).

2.	Song, B.-M., Faumont, S., Lockery, S. & Avery, L. Recognition of familiar food activates feeding via an endocrine serotonin signal in *Caenorhabditis elegans*. *Elife* **2,** e00329 (2013).

3.	Avery, L. & Shtonda, B. B. Food transport in the *C. elegans* pharynx. *J. Exp. Biol.* **206,** 2441–2457 (2003).

4.	Zhang, Y., Lu, H. & Bargmann, C. I. Pathogenic bacteria induce aversive olfactory learning in *Caenorhabditis elegans.* *Nature* **438,** 179–184 (2005).
